# Functional Insights into Dispensable Genes using Genome-Wide Loss-of-Function Burden Tests in Arabidopsis

**DOI:** 10.1101/2025.04.30.651390

**Authors:** Kehan Zhao, Mariele Lensink, J. Grey Monroe

**Affiliations:** Department of Plant Sciences, University of California Davis, Davis CA 95616

## Abstract

Not all genes are essential for plant survival. With the rise of pan-genomics, it is evident that certain genes can be lost without negatively affecting fitness. Naturally occurring loss-of-function (LoF) mutations provide a valuable perspective on gene dispensability, offering insights into deleterious and adaptive gene loss. In this study, we identified 91,751 naturally occurring LoF variants from publicly available Arabidopsis genome data. Our findings demonstrate that LoF-intolerant genes are enriched in essential biological functions and associated with specific histone marks linked to active transcription. In contrast, LoF-tolerant genes exhibit relaxed selective pressure and are enriched in functions related to pollen rejection and defense responses, and can be used as a proxy for dispensable genes in the pan-genome. Using a random forest model trained on histone marks, we achieved moderate success in predicting gene LoF tolerance, with an AUC of 0.717 in Arabidopsis and 0.768 in rice, and even across species. We also pioneered genome-wide LoF burden tests in Arabidopsis, collapsing independent LoF alleles into a single state to reduce allelic heterogeneity. By integrating LoF burden tests with transcriptomic data, we identified thousands of LoF-expression associations. Notably, this analysis accurately recapitulated the flowering time networks and identified *FRIGIDA* as a key regulator of flowering time genes. Furthermore, we found that collapsing alleles based on functional outcomes enhances association sensitivity. These results provide insight into gene dispensability and a framework for leveraging LoF mutations to study gene functions with improved association studies.

## Introduction

Not all genes are essential for organism survival. In the age of pan-genomics, it is clear that many genes within a species’ genome can be absent or lost. Moreover, gene loss can often be adaptive (Smith 1970; Olson 1999; Orr 2005; Murray 2020; Klim et al. 2024). When certain genes become dispensable, their loss can lead to advantageous traits under specific environmental conditions or selective pressures. For example, functional disruption of CBF genes, which encode transcription factors that regulate the expression of various cold-responsive genes, improves fitness in warmer climates in Arabidopsis (Monroe et al. 2016; Lee et al. 2024). In humans, individuals with homozygous knockout for *C-C chemokine receptor type 5* (*CCR5*) are resistant to HIV-1 infection (Liu et al. 1996). A plethora of adaptive gene loss cases has also been reported in other species (Spielmeyer et al. 2002; Wong et al. 2012; Kvitek and Sherlock 2013; Flannick et al. 2014; Gomez et al. 2019; Klim et al. 2024). Adaptive gene loss contributes to the dynamic nature of gene content evolution, where genomes are continuously shaped by gains, losses, and modifications of genes. Previously in human studies, researchers have pointed out that loss-of-function (LoF) variants are valuable for identifying promising drug targets and assessing the safety of gene inhibition (Minikel et al. 2020). Artificial gene knockouts in agriculture, like the knockout of *CYP79D1* and *CYP79D2* in cassava to reduce cyanide levels by 92% (Jørgensen et al. 2005), can accelerate plant breeding and crop improvement.

Studying gene function in the context of dispensability provides insights into which genes are essential for basic cellular processes and which can tolerate LoF mutations. Genes that can withstand LoF are typically non-essential but may play significant roles in specialized or redundant functions. Investigating the functions of these dispensable genes can reveal their contributions to phenotypic diversity and adaptation. In traditional genetic research, the process started by identifying a phenotypic change, followed by discovering the gene responsible for it, often resulting in the identification of new genes. However, genes that had little or no influence on commonly studied traits were largely overlooked. Non-essential genes, by their nature, often don’t have significant effects on observable phenotypes, thus are less studied and have poor functional annotations.

Genome-Wide Association Studies (GWAS) aim to identify genetic variants associated with specific traits by testing for allele-phenotype relationships across large populations (Manolio 2010; Uffelmann et al. 2021). This method aids gene discovery and functional studies by revealing how genes contribute to complex traits and diseases, offering insights into new gene targets. However, allelic heterogeneity poses a significant challenge in conventional GWAS. Allelic heterogeneity occurs when different alleles at a single locus or different loci contribute to the same trait, such as multiple independent LoF variants leading to disrupted gene function, complicating the identification of causal variants (Bergelson and Roux 2010). Over the past decades, the availability of Whole-Genome Sequencing (WGS) data has revolutionized genomic studies, and advances in predicting variant effects from gene sequences have significantly contributed to human and medical research (MacArthur et al. 2012; Brandes et al. 2023; Neville et al. 2024), paving the way for the exploration of genes by leveraging natural genetic variation. To study rare protein-coding variants and enhance the statistical power of their analysis, burden tests were introduced. This approach aggregates genetic variants, typically loss-of-function variants, within a gene to construct a “burden genotype,” which is subsequently tested gene-by-gene for associations with phenotypes. LoF burden tests have proven instrumental in gene discovery within human genomics (Backman et al. 2021; Koprulu et al. 2022; Ivarsdottir et al. 2024; Spence et al. 2024). Although conceptually similar to GWAS, they excel in prioritizing the identification of distinct genes (Spence et al. 2024), offering a complementary and powerful tool for genetic research.

Utilizing LoF alleles for genome-wide association studies offers several additional advantages. Traditional GWAS relies on linkage to genetic markers, which often identify broad genomic regions containing hundreds of candidate genes, necessitating additional work to pinpoint causal loci. By directly associating functional states of individual genes with phenotypes, LoF association testing provides a more targeted approach. Furthermore, LoF burden tests provide actionable insights, offering testable hypotheses for experimental validation.

In this study, we performed genome-wide LoF burden tests that mitigate allelic heterogeneity by collapsing independent LoF alleles to a single allelic state (LoF burden). We tested this approach with transcriptome data, present evidence that collapsing alleles based on their functional outcomes enhances the sensitivity of association testing, and offer a framework for gene function discovery through LoF-expression associations.

## Materials and Methods

### Arabidopsis LoF Calling and LoF Matrix

Loss-of-function variants were called from publicly available Whole-Genome Sequencing data, including SNP variants and small indel calls for 1,135 Arabidopsis accessions from the 1001 Genomes Project (1001 Genomes Consortium 2016) and structural variants and indel calls for 1,301 accessions (Göktay et al. 2021). The VCF from the 1001 Genomes Project annotated using SnpEff was downloaded from 1001 Genomes Data Center (https://1001genomes.org/data/GMI-MPI/releases/v3.1/1001genomes_snpeff_v3.1/). The program snpEff (Cingolani et al. 2012) is widely used to annotate mutation effects from genotype variants. Variants annotated as “High” impact are considered as putative LoF variants. Those include “stop_lost”, “stop_gained”, “start_lost”, “frameshift”, “splice_donor_variants”, and “splice_acceptor_variants”. For structural variants, we mainly focus on deletions as they constitute the majority of the data. A deletion variant is considered a LoF mutation if it either causes a frameshift by disrupting the reading frame (i.e., the deletion overlaps the coding DNA sequence in a length not divisible by three) or removes more than 10% of the coding sequence of the affected transcript. It is also assumed that the protein product could still be functional with a change in the first and last 5% of the coding sequence as an alternative start/stop codon may rescue the disrupted transcript, and thus variants that only affect the first and last 5% of the coding sequence were therefore systematically removed (MacArthur et al. 2012). Lastly, to ensure true knockout of a gene and reduce false positives, only variants that disrupt all gene models for the gene of interest are kept. With the prediction of LoF alleles, a LoF allele matrix is generated to reflect the relationship between accessions and LoF alleles. Independent LoF alleles of the same gene were then collapsed into a single allele state. The collapsed LoF allele matrix represents a binary state of LoF or functional for each accession-gene combination, where LoF alleles are shown as “1” and functional alleles are shown as “0” in the LoF allele matrix (**Supplemental Table 1**).

### Arabidopsis Expression Matrix

An expression matrix was generated based on the 1001 Arabidopsis transcriptomes (Kawakatsu et al. 2016) after correction for batch effect. It reflects the log_2_-transformed transcript level for every accession-gene combination. The NaN values after log_2_ transformation were replaced with the lowest transcript value among all accessions for each gene.

To test if the non-normality of expression data changes the scale of LoF-expression associations, we applied a rank-based inverse normal transformation (INT) to normalize non-normally distributed expression data. Non-missing values were ranked (with ties assigned the average rank) and transformed into a standard normal distribution using the inverse cumulative distribution function of the normal distribution. The transformed values retained the original rank order.

### Gene Ontology Analysis

GO enrichment of LoF-tolerant and LoF-intolerant genes was performed using ShinyGO 0.81 (http://bioinformatics.sdstate.edu/go/) (Ge et al. 2020). An FDR-adjusted p-value cutoff of 0.05 was adopted to identify significantly enriched GO terms.

### Arabidopsis Gene Features and Epigenomic Data

We compiled and generated data on genome-wide sequence and epigenomic features of Arabidopsis to construct a high-resolution predictive model of pan-genome gene dispensability. Polymorphism was previously calculated from the 1001 genomes project (1001 Genomes Consortium 2016). Tissue breadth data were obtained from Mergner et al. (2020). Additionally, we retrieved 62 BigWig-formatted datasets from the Plant Chromatin State Database, detailing the distribution of histone modifications, including H3K4me2, H3K4me1, H3K4me3, H3K27ac, H3K14ac, H3K27me1, H3K36ac, H3K36me3, H3K56ac, H3K9ac, H3K9me1, H3K9me2, and H3K23ac. For each modification, signal intensities were normalized (0 to 1) and averaged across each genomic region for downstream analyses.

### Rice LoF-Tolerant Genes and Epigenomic Data

To obtain a list of LoF-tolerant genes in rice, a set of 29 million biallelic SNPs with SnpEff annotation was downloaded from International Rice Research Institute (IRRI) Rice SNP-Seek Database (https://s3.amazonaws.com/3kricegenome/snpseek-dl/NB_bialSNP_pseudo_canonical_ALL.vcf.gz). Any genes with variants annotated with “High” impact were extracted. To build predictive models of LoF tolerance, the Arabidopsis and rice epigenomic data were obtained from Lu et al. (2019).

### Random Forest Predictions

Random Forest Predictions in this study were performed using the R package “randomForest”. Pan-genome dispensability and gene LoF tolerance were converted to binary factors. The dataset used for model building consisted of annotated genes, with indispensable genes balanced against dispensable genes (or LoF genes balanced against non-LoF genes) through subsampling to ensure equal representation. The predictors (gene features or histone marks) were fitted into the model with 500 decision trees (ntree = 500). Model performance was assessed using receiver operating characteristic (ROC) curves and the area under the curve (AUC), with feature importance quantified by the Mean Decrease in Gini.

### Gene Coexpression Network and Coefficient

To measure gene co-expression, we used previously published data from 874 transcriptomes of natural Arabidopsis genotypes, capturing stochastic interindividual variation. We calculated all-by-all expression correlations using Spearman correlation coefficients, employing the mr2mods program (Kawakatsu et al. 2016; Wisecaver et al. 2017). This data was previously published by Katz et al. (2022).

### Genome-Wide LoF Burden Tests and Permutations

Genome-wide association testing through LoF burdens was performed using EMMAX algorithm (Kang et al. 2010) with the function “cGWAS.emmax” in the R package “cpgen”. The collapsed or uncollapsed binary LoF allele matrix was input as the marker matrix and the expression level of each gene from the expression matrix was input as phenotype for each association testing. The kinship matrix derived from SNP data (https://1001genomes.org/data/GMI-MPI/releases/v3.1/SNP_matrix_imputed_hdf5/) was used in the model to control for population structure.

To mitigate the bias of allele frequency in association studies, we performed 101 permutations for the collapsed and uncollapsed LoF allele matrices, respectively. The order of LoF or functional alleles (“1” or “0” in LoF allele matrix) was shuffled for each gene (or allele for the uncollapsed matrix) but the same allele frequency was maintained. The 101 simulated LoF matrices were input in the LoF association testing pipeline as described above. The p-values from the LoF-expression associations were then compared to and ranked among the simulated values from 100 permutations. A rank matrix was obtained, showing the rank of p-value or the absolute value of beta coefficient among 100 simulations for each LoF-expression association. The 101st simulated p-value matrix was also ranked among the 100 simulations to create null rank matrices.

### Linear Regression Model

Loss-of-function burden tests were also conducted using a linear regression model implemented with the “lm” function in R. Specifically, we modeled gene expression as a function of LoF status.

### Candidate Filtering

To minimize false positives, association hits underwent a series of stringent filtering steps. First, the p-values of LoF-expression associations were corrected for multiple comparisons using the false discovery rate (FDR) method implemented in the “p.adjust” function. Associations with an FDR-adjusted p-value greater than 0.05 were excluded. Second, to eliminate potential linkage effects, gene pairs located within 1,000 kb of each other were removed. Finally, only candidates with the top-ranked (rank 1) p-value among 100 simulations were retained, ensuring only the most robust associations were considered for downstream analysis. While other high-ranking associations were not included in subsequent analysis, they may still represent biologically meaningful signals.

### Gene Network

We constructed and visualized a gene interaction network based on LoF burden-expression associations using the R package “igraph”. Genes are represented as nodes, with edges indicating the relationships between LoF genes and their associated expression genes. The strength of these associations is weighted by statistical significance, while directionality is distinguished based on whether the association is activating or repressing.

## Results & Discussion

### Pan-genome Gene Dispensability Can Be Inferred from Predicted LoF Based on A Single Reference Genome

In total, we called 91,751 naturally occurring variants that have putative LoF effects on genes from publicly available WGS data. This is higher than previous estimates (Xu et al. 2019) because we combined both small scale mutations and structural variants (see ***Materials and Methods***). We emphasize that these are putative loss-of-function similar to related analyses in humans (MacArthur et al. 2012; Emdin et al. 2018; Rausell et al. 2020; Beaumont et al. 2024), and that their true effects would require further experimental validation. The majority of those LoF alleles have lower expression level compared to their functional counterparts, as expected (**Figure S1**). The variant calling process is based on a single reference genome Columbia (Col-0). To evaluate how well LoF calls from a single reference genome translate to pan-genome classifications of gene presence-absence, we applied a machine learning method and trained a random forest model on gene features to predict gene dispensability in the pan-genome. A matrix of gene features, including LoF frequency, tissue breadth, epigenomic patterns, and sequence composition, was used as input predictors. The binary status of gene dispensability was extracted from a recent Arabidopsis pan-genome study of 32 genotypes (Kang et al. 2023), where core genes and soft-core genes were classified as “indispensable” whereas dispensable genes and private genes were classified as “dispensable” in the random forest model. We used 10-fold cross validation for testing the model’s accuracy. Using gene features to predict gene dispensability in the Arabidopsis pan-genome, our random forest model achieves an AUC-ROC of 0.854 (**Fig 1a**), demonstrating strong predictive power of dispensability in the Arabidopsis pan-genome. Among all the predictors, LoF frequency has the highest Mean Decrease Gini score, followed by tissue breadth and H3K4me1. The accuracy and importance of LoF frequency in the predictive model demonstrate the utility of integrating diverse gene features to classify gene dispensability, providing a robust framework to use LoF-tolerant genes inferred from both a single reference genome and small and structural variant calls from a large population genomic dataset as a proxy for dispensable genes in the pan-genome.

**Figure 1.**
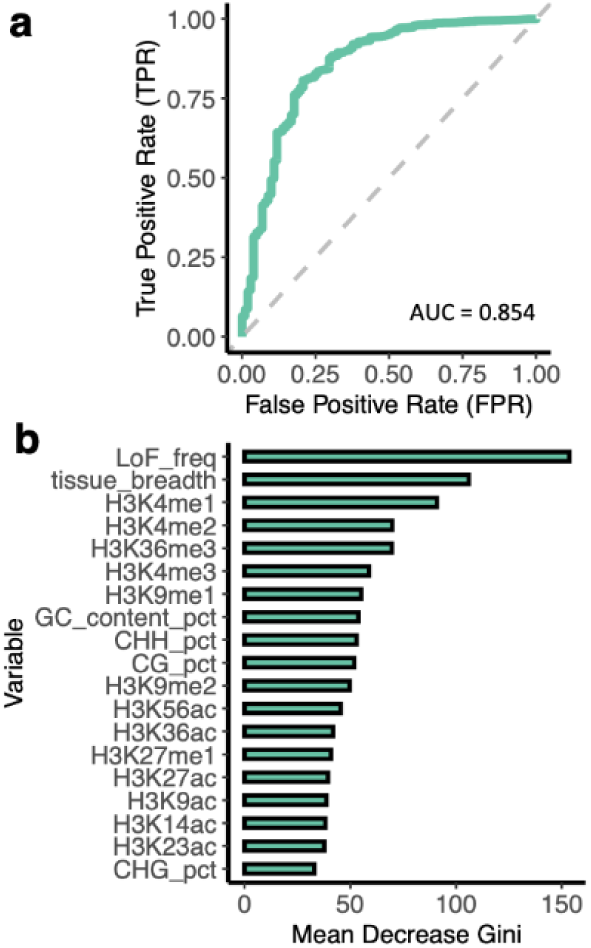
Random Forest Model of Pan-genome Gene Dispensability Prediction. a) The ROC curve of the predictive model using Random Forest (RF). An AUC-ROC value of −0.5 is equivalent to random guessing, while an AUC-ROC of **1** indicates perfect predictions. The diagonal dashed line shows the expected performance of a model based on random guessing. Curves closer to the upper left corner of the chart represent a better predictive performance than curves that are closer to the diagonal dashed line, b) The Mean Decrease Gini of all the predictors in the RF model. Higher Gini score signifies higher importance of the variable to the predictive model.

### LoF-Tolerant and LoF-Intolerant Genes Display Distinct Features in Arabidopsis

Most genes have low LoF frequency across all accessions, and 10,999 genes remain functional (LoF frequency = 0) in all 1,113 accessions represented in the LoF allele matrix (**Figure 2a**). To better understand the distribution of LoF mutations, we categorized genes by their mutational burden: 10,873 genes (39.7%) carry no predicted LoF mutations in their gene body, 5,278 genes (19.3%) harbor only one, and the remaining 41.0% contain multiple independent LoF mutations (**Figure 2b**). Five genes have more than 100 independent LoF alleles accumulating in the natural population. When considering the mutations that lead to loss of function, the most prevalent types of LoF variants we detected were frameshift mutations and premature stop codons, which represent 44.59% and 24.98% of all variants, respectively (**Figure 2c**), consistent with previous reports (Xu et al. 2019).

**Figure 2.**
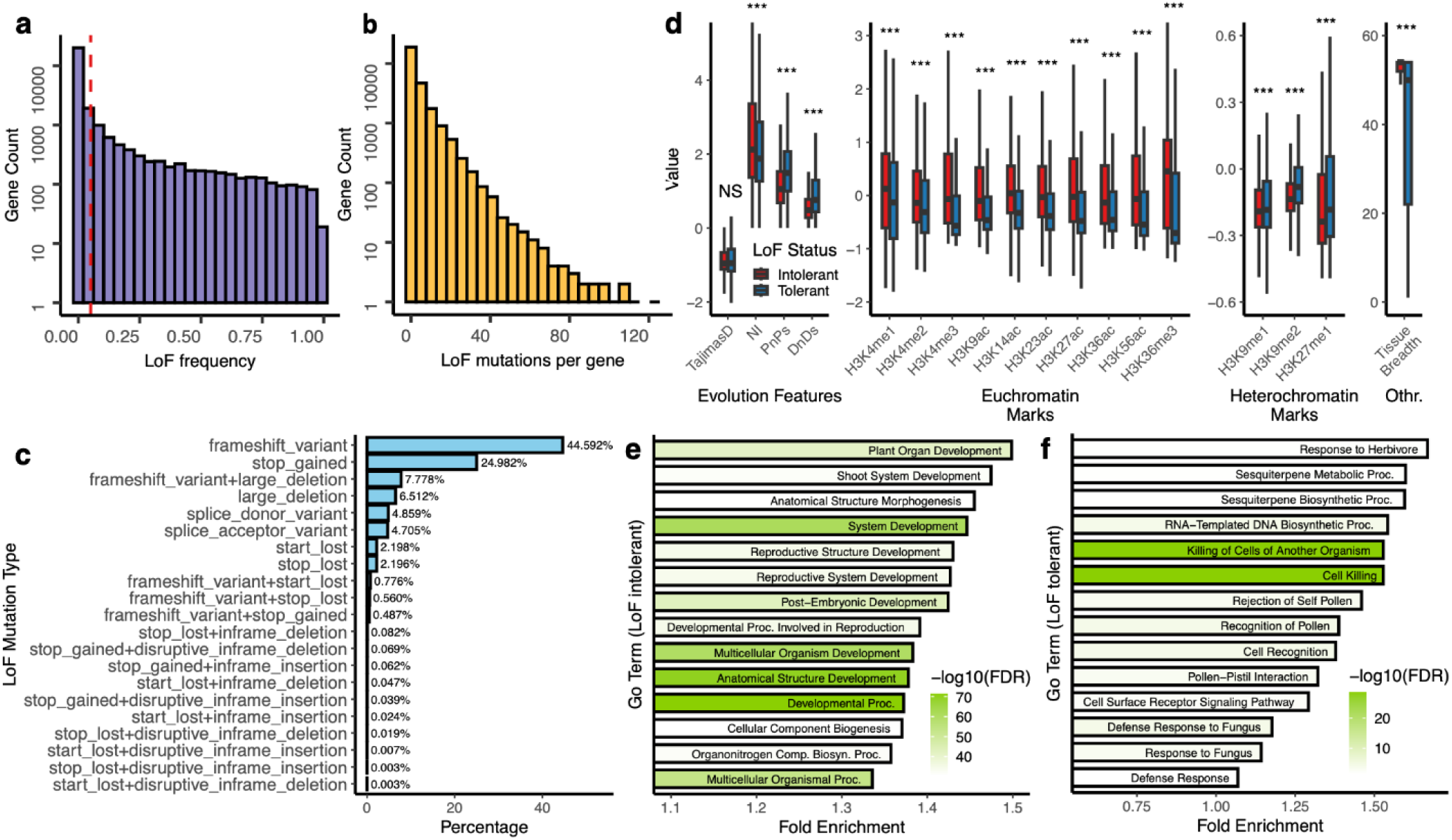
Characteristics of Loss-of-Function (LoF) Mutations in Arabidopsis. a) The histogram of LoF frequency of **all** Arabidopsis genes. Red dash line indicates the 5% allele frequency threshold commonly used in Genome-Wide Association Studies, b) The histogram of numbers of independent LoF mutations per gene, c) Percentage of each mutation type that causes gene disruption, d) Box plots of gene features of LoF-tolerant and LoF-intolerant genes in Arabidopsis. e) Top 14 Gene Ontology (GO) terms of LoF-intolerant genes in Arabidopsis ranked by Fold Enrichment, f) Significant Gene Ontology (GO) terms of LoF-tolerant genes in Arabidopsis ranked by Fold Enrichment.

Chromatin states, such as histone modifications, are closely linked to gene regulation and function (Berr et al. 2011; Deal and Henikoff 2011; Liu et al. 2018). Because these epigenomic features can reflect the functional importance of genes, we investigated whether gene tolerance to loss of function is associated with specific chromatin marks. Genes were categorized into two groups according to their tolerance to loss of function (tolerant: LoF frequency > 0). We found that genes intolerant to LoF are enriched in H3K4me1, H3K4me2, H3K4me3, H3K9ac, H3K14ac, H3K23ac, H3K27ac, H3K36ac, H3K36me3, and H3K56ac (**Figure 2d**). These epigenomic marks are generally associated with active gene transcription (Roth et al. 2001; Bannister and Kouzarides 2011; Deal and Henikoff 2011; Liu et al. 2018; Lee et al. 2020; Jamge et al. 2023). However, the same enrichment was not observed in H3K27me1, H3K9me1 and H3K9me2, which have been previously shown linked to gene repression or heterochromatin formation (Noma K et al. 2001; Schotta et al. 2002; Deal and Henikoff 2011; Pan et al. 2018). These findings are similar to that in humans where loss-of-function intolerant genes are enriched for active epigenomic marks (Boukas et al. 2022). We further observed higher Pn/Ps, higher Dn/Ds ratios and lower neutrality index (NI) in genes tolerant to LoF, consistent with these genes experiencing less purifying selection and reduced functional constraint compared to genes intolerant to LoF. LoF-intolerant genes are also expressed in a wider range of tissue types (higher tissue breadth), indicating their roles across diverse biological processes and developmental stages. Interestingly, we also found that genes highly expressed in reproductive tissues exhibit significantly lower LoF rates compared to the genome-wide average (**Figure S2a**). This underscores the critical importance of reproduction-related genes, which appear to be under stronger selective pressure. As expected, genes with prominent mutant phenotypes (Lloyd and Meinke 2012) also exhibit lower LoF frequencies compared to other genes (**Figure S2b**).

Moreover, we performed functional analysis of LoF-tolerant and LoF-intolerant genes using Gene Ontology (GO) terms. LoF-intolerant genes are enriched with functionally important GO terms that are vital to plants’ growth, reproduction, and development (**Figure 2e**). This is expected as detrimental LoF mutations in functionally important genes would have been purged by purifying selection. Conversely, the GO terms for LoF-tolerant genes (LoF frequency > 0) are enriched in sesquiterpene metabolic process and defense-related GO terms, including “Killing of Cells of Another Organism” and “Defense Response” (**Figure 2f**). Sesquiterpenes are a diverse group of naturally occurring terpenes and play crucial roles in plants, including defense mechanisms, environmental stress responses, communication and signaling, etc (Chizzola 2013). Plants maintain a delicate balance in allocating energy and resources between defense and growth (Giolai and Laine 2024). Previous research found that a mutation in *Replication Factor C Subunit 3* (*RFC3*) enhances resistance to pathogen infection but also leads to developmental defects, including smaller plant size, narrower leaves and petals, and reduced cell proliferation (Xia et al. 2009). In certain environments, it may be advantageous for plants to lose genes associated with defense mechanisms, enabling greater allocation of resources toward overall growth and reproduction. The regulation of biotic and abiotic stress responses is highly interconnected, with extensive signaling cross talk shaping plant adaptation (Jambunathan et al. 2001; Paparella et al. 2014; Desaint et al. 2024). Recent studies highlight this trade-off: a disease resistance gene in wild pepper was repeatedly lost at high temperatures, indicating that pathogen resistance may entail fitness costs (Poulicard et al. 2024), while species with lower pathogen pressure exhibit pronounced loss of immune receptors during adaptation (Li et al. 2025). Moreover, the convergent loss of plant immunity components across multiple lineages underscores the complex coevolutionary interplay between the immune system and drought tolerance, as these genes are differentially regulated in defense and drought response pathways (Baggs et al. 2020). Interestingly, some cases of gene loss enhance immunity without compromising growth and architecture (Castelló et al. 2010; Wang et al. 2019), making these genes particularly valuable for agricultural applications.

LoF-tolerant genes in Arabidopsis are also associated with “Rejection of Self Pollen”, “Recognition of Pollen” and “Pollen-Pistil Interaction”. Because Arabidopsis is a dominantly inbreeding plant species, the predominance of inbreeding may reduce the need for or even select against the function of genes that mediate pollen-pistil interactions and self-incompatibility. Notably, the GO terms associated with LoF-tolerant genes show substantial overlap with those of genes affected by copy number variations (CNVs) as reported by (Zmienko et al. 2020). These terms are significantly enriched for processes related to interactions with other organisms, defense mechanisms, and stress responses. Furthermore, LoF mutations are more commonly tolerated in non-single-copy genes, as expected (Xu et al. 2019) (**Figure S2c**).

We also observed a positive correlation between gene CDS length and the number of accumulated LoF variants (p < 2.2 × 10^−16^), which appears to depend on the functional importance of the gene. However, despite this overall trend, some genes accumulate relatively few LoF variants regardless of their length. This may indicate that these genes are under stronger selective pressures, possibly due to their constrained biological functions. (**Figure S3**).

### Within- and Cross-Species Prediction of LoF Tolerance through Epigenomic Patterns

We next set out to test if epigenomic information could serve as a broadly applicable predictor of gene dispensability in plants. We focused on histone modifications which reflect gene activation and repression, to predict LoF tolerance of genes. Using a random forest model trained with *Arabidopsis thaliana* histone marks, we achieved a moderate predictive performance for gene LoF tolerance within the species, with an AUC-ROC of 0.717 and H3K4me3 being the top predictor (**Figure 3a&b**). We applied the same approach to rice (*Oryza sativa*) by training a separate random forest model with its corresponding histone marks, achieving a similar result with an AUC-ROC of 0.768 and H3K4me3 being the most important predictor (**Figure 3c&d**). These results indicate that epigenomic features can be used to infer LoF tolerance. To further evaluate the generalizability of these models, we tested them across species. When the Arabidopsis-trained model was applied to rice data, it achieved an AUC-ROC of 0.724 (**Figure 3e**), comparable to within-species predictions for Arabidopsis. Conversely, the rice-trained model tested on Arabidopsis data yielded a lower AUC-ROC of 0.606 but still outperformed a random expectation (**Figure 3f**). This cross-species validation underscores the potential of conserved epigenomic features as reliable predictors of gene LoF tolerance across diverse plant species.

**Figure 3.**
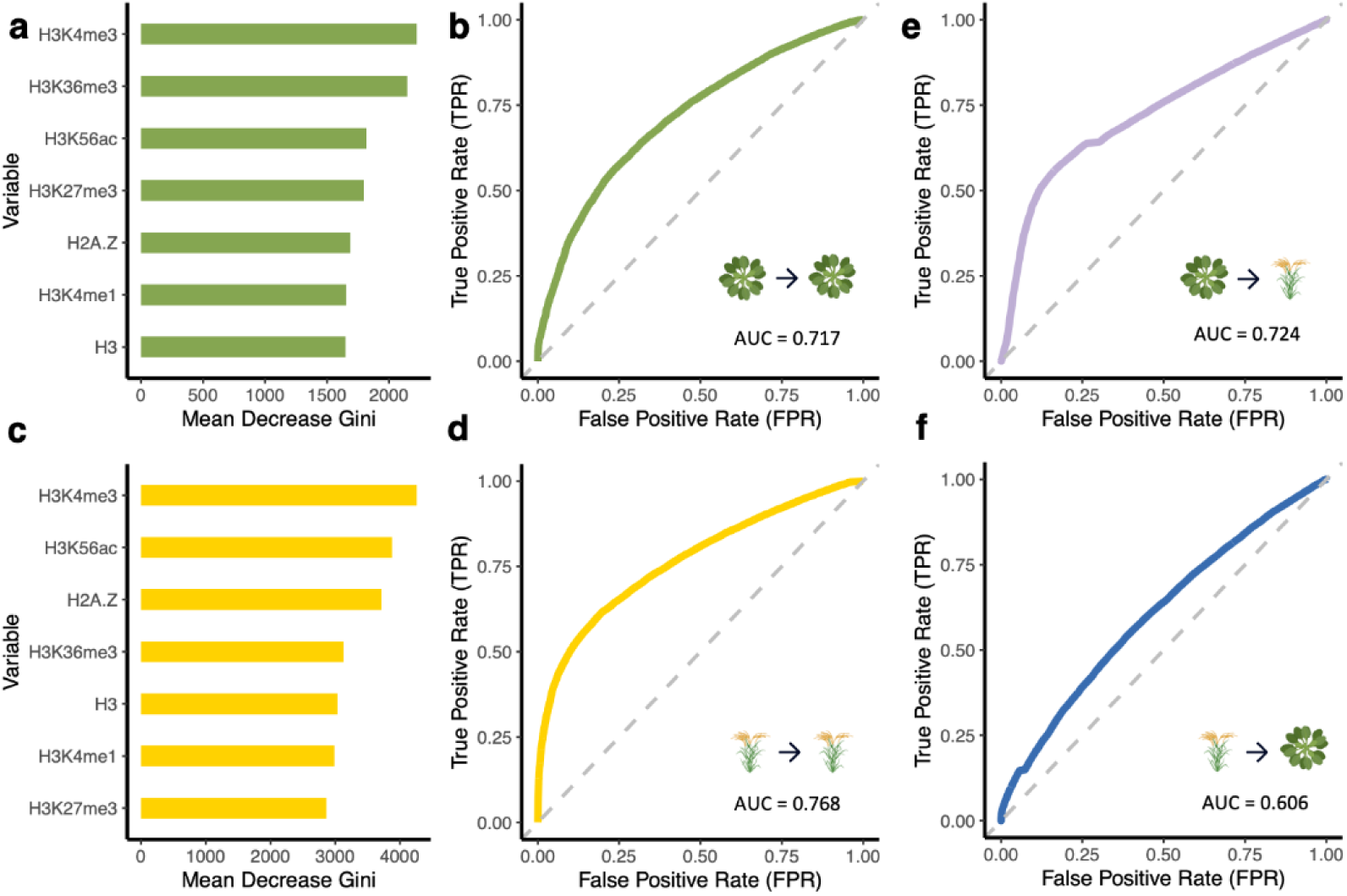
Within- and Cross-Species Prediction of LoF Tolerance through Epigenomic Patterns. a) The Mean Decrease Gini of histone marks in predicting Arabidopsis LoF tolerance, b) The ROC curve of the random forest model trained on Arabidopsis thaliana histone marks to predict Arabidopsis LoF tolerance, c) The Mean Decrease Gini of histone marks in predicting rice LoF tolerance, d) The ROC curve of the random forest model trained on rice histone marks to predict rice LoF tolerance, e) The ROC curve of the Arabidopsis-trained random forest model tested on rice data, f) The ROC curve of the rice-trained random forest model tested on Arabidopsis data. For b, d, e & f), the diagonal dashed line shows the expected performance of a model based on random guessing. Curves closer to the upper left corner of the chart represent a better predictive performance than curves that are closer to the diagonal dashed line.

### Collapsing LoF Alleles into Their Functional States Mitigates Allelic Heterogeneity in Association Testing

Gene loss of a locus can be caused by multiple independent molecular variants, for example, a frameshift mutation or the introduction of a premature stop codon. This phenomenon is called allelic heterogeneity and can be problematic for the conventional functionally agnostic genome-wide association studies, potentially masking important loci (Monroe et al. 2021). Categorizing variants by their predicted functional effects should theoretically increase our ability to detect causative loci exhibiting allelic heterogeneity. Building on this idea, loss-of-function burden tests have been developed to aggregate the effects of multiple rare or independent LoF variants within a gene into a single burden score. By collapsing these variants at the gene level, this approach enhances statistical power to detect associations between gene disruption and phenotypic variation – even when individual variants are too rare to show significant effects on their own (Povysil et al. 2019). Based on the prediction of variant effects on gene function, we created uncollapsed and collapsed LoF matrices, respectively (***Materials and methods,* Figure 4a**). Gene expression as a class of quantitative traits can be highly heritable and have complex genetic architecture (Kliebenstein 2009). In this study, we utilized gene expression as a suite of quantitative traits to evaluate the performance of LoF burden tests for gene discovery in association studies in plants, testing for the effects of candidate loci on the expression of thousands of genes at a time. Using the collapsed LoF matrix and transcriptome data from the 1001 Genomes collection of *Arabidopsis thaliana* (Kawakatsu et al. 2016), we performed genome-wide LoF burden-expression tests, which yield 14,175 significant associations after stringent filtering processes (***Materials and methods,* Figure 4b**). The loss of function of one gene is associated with the expression of up to 175 other genes, although the majority of LoF genes are associated with the expression of fewer than 50 genes (**Figure 4c**). Similarly, individual expressing genes are associated with up to 80 expression quantitative trait loci (eQTLs), with most genes linked to fewer than 20 eQTLs (**Figure 4d**). Statistically, the power to detect associations increases with the allele frequency of a variant, which is why common genetic variants have been extensively studied, while rare variants remain a compelling frontier in GWAS (Lee et al. 2014; Auer and Lettre 2015). Interestingly, we observed a higher number of significant LoF-expression associations for genes with either very low or very high LoF frequencies (**Figure S4**). A possible explanation for this pattern is that genes with low LoF frequencies are likely to be functionally pleiotropic, and their loss is more likely to affect the expression of numerous other genes. Alternatively, this result could reflect a bias in detection of non-normally distributed phenotypes with rare variants (Beasley et al. 2009; McCaw et al. 2020), which we therefore aimed to address.

**Figure 4.**
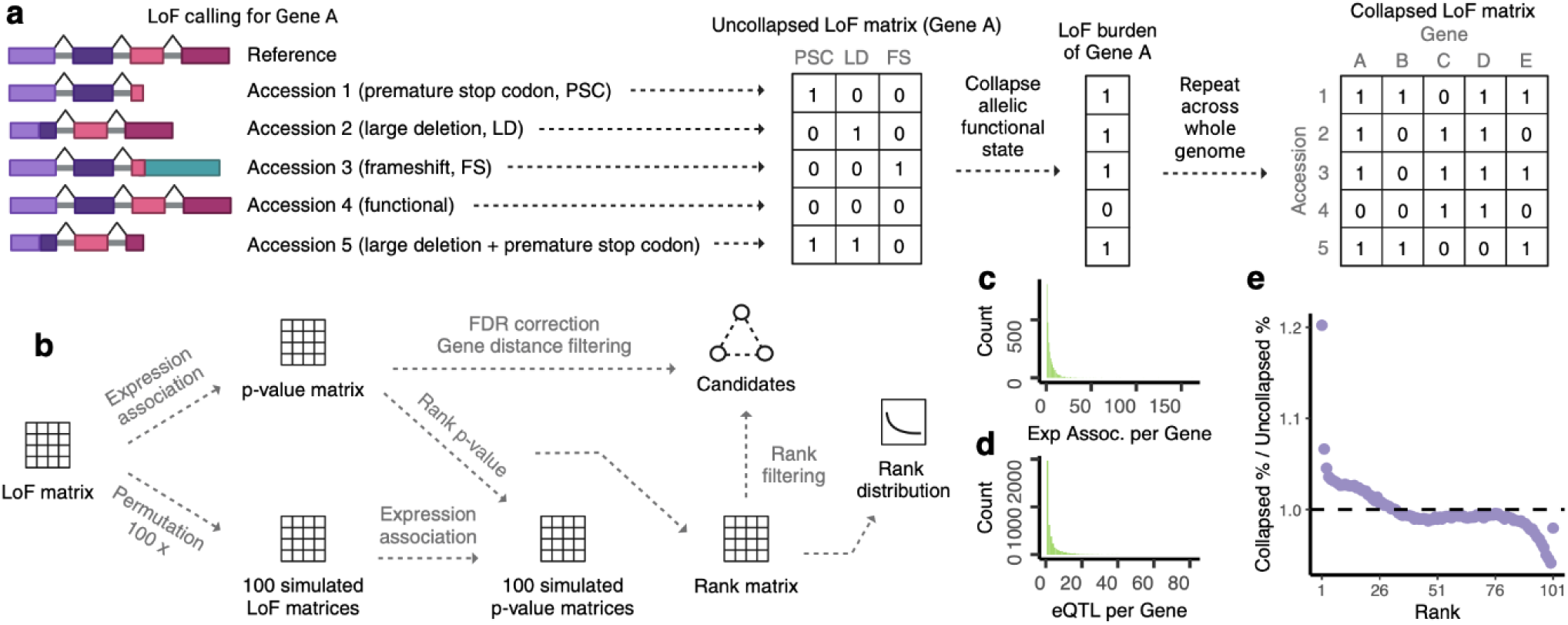
Collapsing LoF Alleles into LoF Burdens Mitigates Allelic Heterogeneity. a) An diagram illustrating LoF calling from a single reference genome and the creation of LoF matrix. The LoF matrix reflects a binary state of LoF or functional for each accession-gene combination, so independent LoF alleles in the species are collapsed into a single allele state, where **1** indicates LoF and 0 indicates a functional allele, b) An illustration of LoF-expression association workflow and candidate filtering process, c) The histogram of number of expression associations per LoF gene, d) The histogram of number of eQTLs per expression gene, e) The ratio of association percentage at different ranks for collapsed and uncollapsed LoF association testing. More high-rank associations were identified with the LoF burden approach. Dash line represents null expectation (ratio = **1).**

To further reduce false positives and eliminate the bias of allele frequency, the p-value of each association testing was ranked among the first 100 simulations, and the 101st simulation was also ranked among the same 100 simulations to generate a null distribution of the rank matrix (**Figure 4b**). An enrichment of high ranks (e.g. rank 1-10) was observed in the LoF-expression associations while the null scenario is evenly distributed (**Figure S5**), suggesting more significant associations between gene LoF and gene expression than expected by chance. To test if collapsing LoF alleles mitigates allelic heterogeneity and as a result, increases the power of association testing, we performed the same approach with the uncollapsed LoF allele matrix. Overall, a reduction in high-rank and an increase in low-rank LoF-expression associations were observed with the uncollapsed LoF allele associations. Specifically, there was a 17.2% decrease in the percentage of rank 1 associations (**Figure 4e, Figure S5**). This aligns with the expectation that LoF burden approaches which contrast alleles based on their functional state rather than individual variants or linked SNPs have improved power by overcoming allelic heterogeneity. Intriguingly, we observed a slight enrichment of rank 101 associations in both collapsed and uncollapsed tests. This pattern is likely attributable to correction for population structure in a GWAS framework, caused by extreme covariance between allele states and the kinship matrix. To further investigate, we performed associations without accounting for kinship, using a simple linear regression model. While this approach resulted in no enrichment at low ranks (e.g. rank 90-101) and a much higher number of rank 1 associations, as expected for true positives, many still are likely false positives driven by spurious correlations due to population structure (**Figure S5**).

### LoF-expression Associations Recapitulate the Canonical Flowering Regulatory Network

In the global analysis of LoF-expression associations, we identified 11,445 significant positive (e.g. LoF in gene A is associated with increased expression in gene B) and 2,730 negative associations for a ratio of positive/negative of 4.2 (**Supplemental Table 2)**. These associations can be visualized as a gene network that captures the significance and directionality of each association (**Figure 5a**). To assess whether the unbalanced ratio of positive to negative associations has biological significance or is merely a statistical artifact, we applied the same filtering process to 50 simulated datasets generated by permuting LoF and functional alleles. Surprisingly, the simulations produced even higher positive-to-negative association ratios (**Figure S6a**). This indicates that the inflation of positive associations could be largely driven by artifacts in the association testing. However, the number of biologically meaningful positive LoF-expression associations we observed is actually lower than expected by chance, suggesting that the real data may be less influenced by statistical noise than the permuted datasets and therefore more reflective of true underlying biological relationships. To explore whether the non-normal distribution of expression data contributes to this bias, we applied a rank-based inverse normal transformation (INT) (McCaw et al. 2020). However, INT did not fully normalize the expression data, as 8,589 genes (37.5% of all expressed genes) remained non-normally distributed due to an abundance of zero-value observations (Shapiro-Wilk normality test, p < 0.05). After excluding these genes, the positive-to-negative association ratio dropped to 0.63, while the 50 simulations averaged 1.06 (One-Sample t-test, p < 2.2 × 10^−16^, **Figure S6b**). These findings suggest that the observed bias toward positive associations is driven by enrichment of type I errors for positive associations due to the skewed distribution of the expression phenotype and unequal group sizes in the binary predictor, which should be carefully considered in future eQTL studies, particularly when analyzing associations with rare alleles. Additionally, the higher prevalence of negative LoF-expression associations suggests that many genes may function to promote the expression of others, consistent with widespread positive regulatory interactions in the transcriptome.

**Figure 5.**
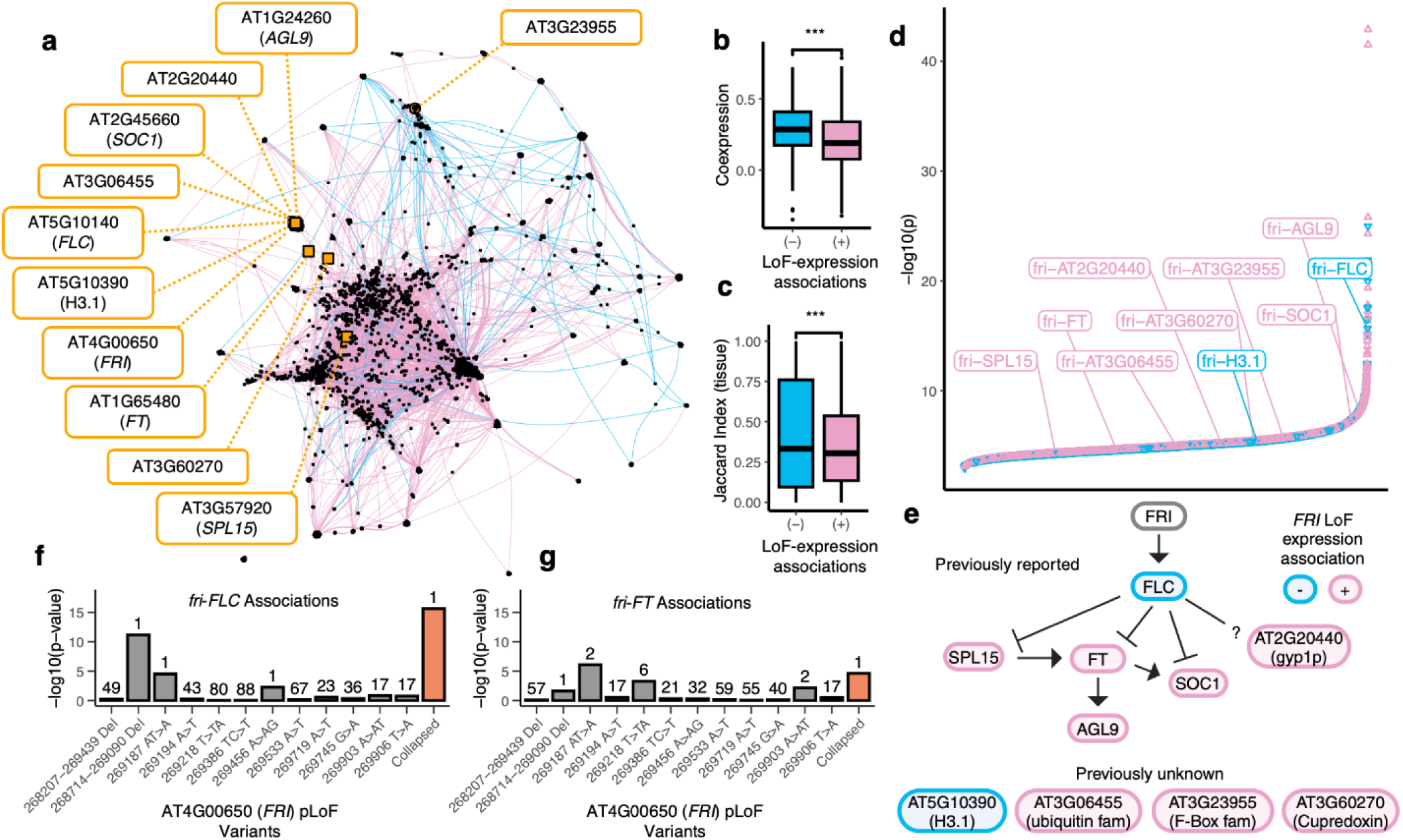
LoF Burden-Expression Association Recapitulates the Canonical Flowering Regulatory Network. a) The gene network constructed from LoF burden-expression association results. Nodes in the network represent genes, categorized as either LoF genes (circles) or Exp genes (squares). Genes serving as both LoF and Exp genes are depicted as circles. The edges denote directed associations from LoF genes to Exp genes, with edge colors indicating the direction of the association: pink for positive associations and blue for negative associations. Flowering time network genes identified by LoF burden-expression association are highlighted with orange nodes and labels, b) Coexpression coefficients of negative and positive LoF-expression association gene pairs, c) Jaccard Index of tissue-specific expression profiles of negative and positive LoF-expression association gene pairs. Higher Jaccard Index means greater similarity between the tissue-specific expression profiles of two genes, d) LoF-expression associations ranked by -log10(ρ). Positive (pink) and negative (blue) associations are distinguished, with *FRI* associations labeled, e) The flowering regulatory network described in previous studies and signified by LoF burden-expression association. Arrows indicate gene activation, and blunted lines indicate repression. Pink: positive expression associations with *FRI* LoF; Blue: negative expression associations with *FRI* LoF. f & g) The uncollapsed and collapsed association results for *fri-FLC* and *fri-FT* associations, respectively. Bars indicate uncollapsed (gray) and collapsed (orange) association significance, with x-axis labels showing variant positions or deletions. Numbers above bars indicate the rank of LoF-expression associations across 100 simulations.

Given the concerns about the imbalance between positive and negative associations, we next asked whether the directionality of these associations aligns with known gene functional relationships inferred from independent datasets. Subsequently, we constructed gene coexpression networks using Spearman correlation based on gene expression data. The global patterns revealed that gene pairs with negative LoF-expression Beta coefficients exhibit higher coexpression coefficients compared to those with positive Beta coefficients (**Figure 5b**). To explore the spatial linkages of these significant gene pairs, we calculated the Jaccard similarity index of their tissue-specific expression profiles, finding that gene pairs with negative Beta coefficients also tend to express in similar tissues (**Figure 5c**). A gene pair with a negative LoF-expression Beta coefficient is positively co-regulated – perhaps part of the same pathway or functional module, where the activity of one gene supports or enhances the activity of the other (Eisen et al. 1998). It is natural that they are also expressed in similar tissues as genes involved in the same biological processes often exhibit tissue-specific coexpression because their function is required in specific cell types or environmental conditions (Marbach et al. 2016; Sonawane et al. 2017). On the other hand, genes in antagonistic relationships are less likely to be directly coexpressed because their regulation is inverse rather than synchronous. Consequently, lower coexpression coefficients are expected for gene pairs with positive Beta coefficients. These findings underscore a strong functional and spatial connection between significantly associated LoF-expression gene pairs, suggesting their potential involvement in coordinated biological processes.

Due to the poor annotation of many less-studied non-essential genes, interpreting many of these associations remains challenging. Nevertheless, the LoF burden-expression tests effectively captured associations between *FRIGIDA* (*FRI*), a major determinant of natural variation in Arabidopsis flowering time, and its downstream targets in the well-studied flowering time pathway. These associations exhibit remarkable statistical significance, with directionalities that align with findings from previous empirical studies (**Figure 5d&e**). Specifically, the LoF of *FRI* is negatively correlated with the expression of *FLOWERING LOCUS C* (*FLC*), consistent with earlier reports that *FRI* activates *FLC* transcription (Clarke and Dean 1994; Gazzani et al. 2003; Choi et al. 2011). Additionally, positive associations were observed between the LoF of *FRI* and the expression of *FLOWERING LOCUS T* (*FT*) and *SUPPRESSOR OF OVEREXPRESSION OF CONSTANS1* (*SOC1*), which is consistent with findings that *FT* facilitates the activation of *SOC1* (Yoo et al. 2005) and that *FLC* binds and repress *FT* and *SOC1* (Deng et al. 2011).

LoF burden-expression tests also identified a positive association between *FRI* LoF and *AGAMOUS-LIKE 9* (*AGL9*) expression, a gene downstream of *FT* that promotes flowering (Mandel and Yanofsky 1998; Yu and Goh 2000; Hsu et al. 2003; Gao et al. 2021). Similarly, a positive association was noted with *SQUAMOSA PROMOTER BINDING PROTEIN-LIKE 15* (*SPL15*), known to promote *FT* (Kim et al. 2012), mediate floral transition (Xu et al. 2016), and which is repressed by *FLC* (Deng et al. 2011). Another notable association is with the expression of AT2G20440, a Ypt/Rab-GAP domain protein whose knockout results in early flowering (Chien et al. 2023), though the molecular mechanism and its interactions with other flowering genes warrant further investigation.

Furthermore, our results uncovered novel associations involving *FRI*, including genes such as AT5G10390 (HISTONE 3.1), AT3G06455 (ubiquitin family protein), AT3G23955 (F-box family protein), and AT3G60270 (Cupredoxin superfamily protein). Notably, LoF burden-expression tests not only detected direct targets of the LoF gene but also downstream pathway components, exemplified by capturing associations between *FRI* and *FT*, with *FLC* as an intermediary (**Figure 5d**). These findings highlight the power of LoF burden-expression tests to dissect complex regulatory networks, offering novel insights into gene interactions and uncovering both known and previously uncharacterized biological connections.

Because the *fri*-*FLC* relationship serves as a positive control whose causality is extensively supported experimentally, it served as a test case for examining the consequence of allelic heterogeneity in gene discovery. We therefore compared the uncollapsed LoF-expression association results to the collapsed LoF burden approach, focusing on associations involving *FRI*. When associating with *FLC* expression, collapsing LoF alleles resulted in a notably reduced p-value, underscoring the increased sensitivity of the collapsed approach. Interestingly, the strongest signal in the uncollapsed association testing was driven by a large structural deletion (**Figure 5f**). This finding is particularly striking, as structural variations have often been underexplored in traditional GWAS analyses. In the case of *FT* association, the collapsed LoF burden test produced a slightly less significant p-value compared to the most significant uncollapsed variant. However, the LoF burden approach achieved a higher rank across simulations (**Figure 5g**), suggesting greater robustness in identifying biologically relevant associations. Overall, these observations highlight that collapsing alleles with shared functional consequences may reduce noise and better capture the cumulative effects of LoF variants on trait variation. This further supports the utility of collapsing LoF alleles in LoF burden tests as a strategy to uncover subtle but meaningful genetic associations, particularly in traits influenced by complex genetic architectures.

### LoF Burden Tests with Flowering Time Capture *FRIGIDA* as a Key Regulator

Inspired by the identification of the flowering time network through LoF burden tests with gene expression data, we sought to investigate whether association tests with phenotypic traits, such as flowering time, could yield similar insights. To explore this, we conducted genome-wide LoF burden tests for flowering time at 10 °C and 16 °C using EMMAX. As anticipated, *FRI* LoF emerged as a highly significant gene for flowering time at both temperatures, with results nearing or surpassing a stringent Bonferroni-corrected p-value threshold. However, no other significant associations were identified under these conditions (**Figure 6a&c**).

**Figure 6.**
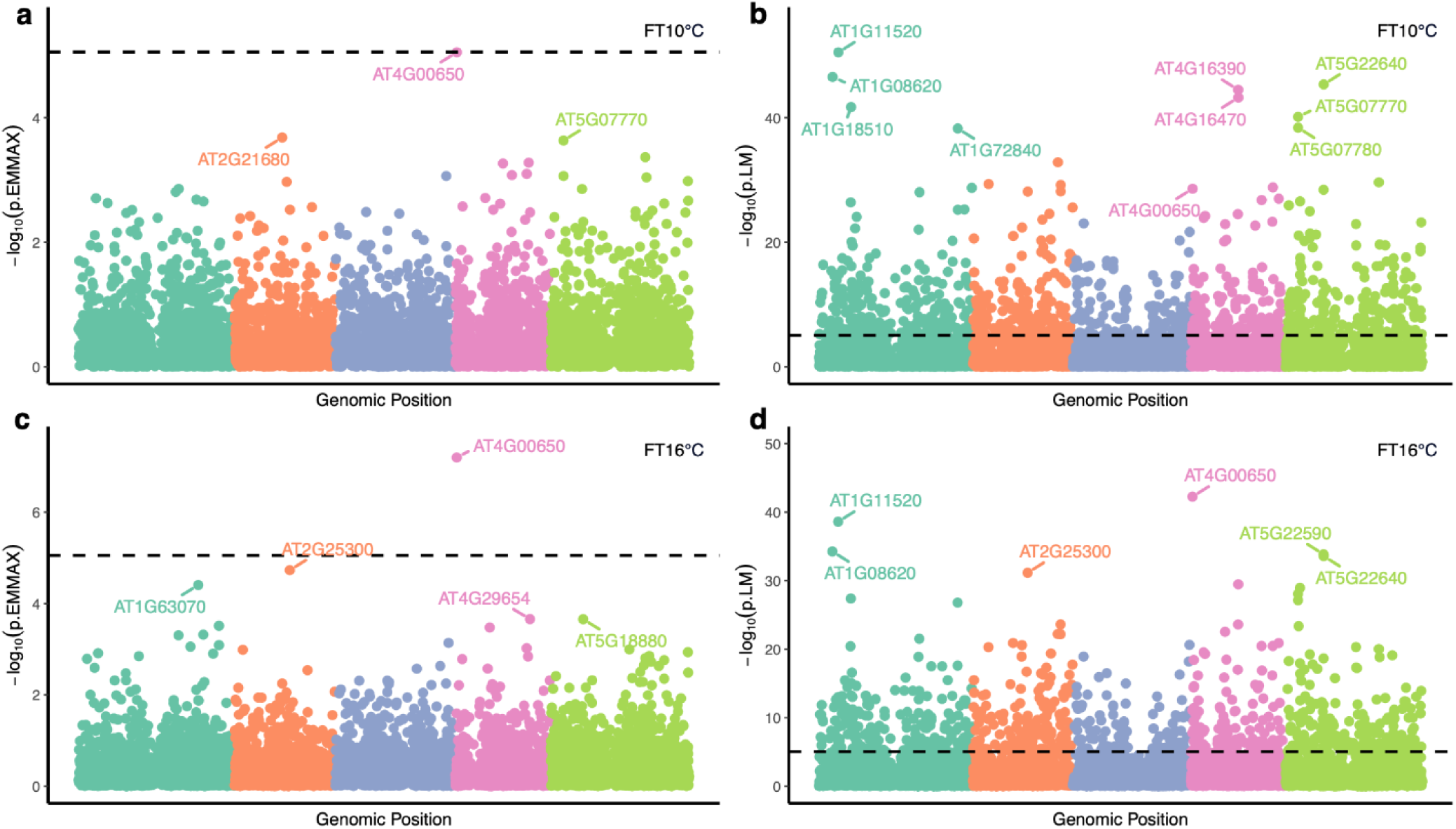
LoF Burden Tests with Flowering Time Capture *FRIGIDA* as a Key Regulator. a & c) Manhattan plots showing genome­wide LoF burden tests using EMMAX with flowering time at 10°C and 16°C, respectively. Each point represents the LoF burden test for a specific gene, with colors indicating the chromosome on which the gene is located. The genomic position of each gene is determined as the midpoint between its start and end positions, based on the Col-0 gene annotation. The dashed line indicates the significance threshold determined by Bonferroni correction, with *FRIGIDA* (AT4G00650) being the only gene surpassing this threshold, b & d) Manhattan plots showing genome-wide LoF burden tests using a linear model approach with flowering time at 10°C and 16°C, respectively. Significantly more associations surpass the Bonferroni threshold due to the lack of population structure control.

Because correction for population structure can obfuscate true causal relationships, we also performed a simplified version of the LoF burden tests using a simple linear model. Under this less stringent approach, a considerable number of associations surpassed the Bonferroni threshold, in addition to *FRI* (**Figure 6b&d**). Distinguishing whether these signals represent false positives due to uncorrected population stratification or if they are genuine, biologically meaningful associations is challenging. A notable case in this regard is AT1G11520, which appeared among the most significant hits at both 10 °C and 16 °C. Previous research by Liu et al. (2021) used a deletion-based genome-wide association approach and found that accessions carrying a 182-bp deletion in AT1G11520 exhibited delayed flowering. Interestingly, all accessions from northern Sweden contained this deletion, highlighting a strong population structure. This high degree of population stratification likely explains why this signal was lost when population structure was controlled for in our analysis. Other highly significant associations revealed here also yield compelling candidates (**Supplemental Table 3**). For example, multiple MADS-box genes show associations with flowering time at 10 °C, including AT2G41445 (p = 6.43 × 10^−30^), AT2G41440 (p = 6.71 × 10^−29^), AT4G11880 (p = 9.69 × 10^−12^), and AT2G41470 (p = 1.84 × 10^−9^), etc. Members of the MADS-box gene family are well known for their roles in the transition to flowering (Parenicová et al. 2003). In particular, AT4G11880 (*AGAMOUS-LIKE 14*, *AGL14*) has previously been shown to promote flowering (Pérez-Ruiz et al. 2015; Chen et al. 2022), aligning with our finding that loss-of-function of AGL14 is positively associated with flowering time, despite its low LoF allele frequency (LoF frequency = 0.036).

Similarly, loss of function in AT1G08620 (*JUMONJI DOMAIN-CONTAINING PROTEIN 16*, *JMJ16*) is significantly and positively associated with flowering time at both 10 °C and 16 °C, suggesting a role for *JMJ16* in promoting flowering. The Arabidopsis genome encodes six homologs of Lysine Demethylase 5 (KDM5): *JMJ14*, *JMJ15*, *JMJ16*, *JMJ17*, *JMJ18*, and *JMJ19* (Lu et al. 2008). While previous studies reported no discernible phenotype for *JMJ16* mutants (Yang et al. 2010), other members of this family – such as JMJ14, which represses the floral transition (Yang et al. 2010), and *JMJ15* and *JMJ18*, which promote flowering (Yang et al. 2012a, 2012b) – support the plausibility of a regulatory role for *JMJ16* as well. We should also emphasize that although we cannot fully exclude the possibility of spurious associations due to limited control for population structure in these cases, the strength of the associations suggests they may reflect true biological effects and thus merit further investigation.

These findings highlight an important trade-off in genetic association studies: while controlling for population structure is necessary to reduce false positives, it also carries the risk of masking true biologically meaningful signals. In cases like AT1G11520, where population structure is intrinsically tied to the phenotype of interest, this trade-off can obscure valuable insights. Thus, while controlling for population structure remains a best practice to mitigate confounding effects, careful consideration must be given to the potential for missing true positive signals when this control is applied.

### Current Limitations and Potential Improvements

Our LoF-calling strategy, while effective, is not without its limitations. Using a single reference genome simplifies analyses but inherently limits our ability to capture variation across diverse genomes. Genes absent in the reference genome or structural rearrangements that deviate from it cannot be accurately assessed. For example, tools like SnpEff rely on genome annotations that, while powerful, cannot perfectly predict the functional impact of all variants. Furthermore, our approach considers genetic variants individually. In rare cases, LoF effects may be rescued by compensatory mutations, such as a frameshift mutation restored by a subsequent deletion, which can lead to functional outcomes.

However, as a famous saying goes “All models are wrong, but some are useful” (Box 1976), our genome-wide LoF burden tests successfully captured the textbook flowering regulatory networks centered around *FRI*, validating its usefulness. Notably, a naturally occurring 16-base-pair deletion is found in the loss-of-function *FRI* allele in Col-0 (Johanson et al. 2000; Risk et al. 2010). This demonstrates that even with imperfect reference-based predictions, functional outcomes of genetic variants can still be inferred. The *FRI* associations, while not the most statistically significant among all LoF-expression association results (**Figure 5d**), exemplifies the power of LoF burden tests to uncover biologically meaningful signals. This suggests that our dataset of LoF-expression associations represents a unique resource which may contain other true biological interactions warranting further exploration (**Supplemental Table 2**).

To address some of the challenges mentioned above, several improvements can enhance LoF prediction in future studies. To begin with, transitioning from single-reference genomes to pan-genomes would enable better representation of genetic diversity across populations, capturing dispensable genes that are absent in the reference. Moreover, integrating tools like AlphaFold could help predict how specific genetic variants impact protein function, providing a more precise definition of LoF.

The rapid development of long-read sequencing technologies and improved assembly algorithms has also highlighted the importance of structural variants in shaping genome function and evolution. Long reads enable more accurate characterization of complex variants, such as insertions, deletions, and rearrangements, that are often missed by short-read technologies (Chaisson et al. 2019; Zhao et al. 2021; Ahsan et al. 2023). As structural variants play a significant role in gene function and dispensability, incorporating this information will be critical in refining LoF predictions. At the same time, LoF studies based on short-read sequencing provide a valuable glimpse into gene dispensability that preempts large pan-genomes, as our findings demonstrate that LoF frequency is the strongest predictor of non-core genes in the pan-genome.

An intrinsic limitation of our methodology is the statistical challenge posed by low allele frequency. In standard GWAS, detecting significant signals is generally easier when the minor allele frequency (MAF) of a variant is higher. Variants with a MAF below 5% are often excluded from analysis to maintain statistical power and minimize false positives. To ensure the robustness of our candidate selection, we adopted this approach in our study. However, this filtering led to the removal of a large proportion of genes (**Figure 2a**), particularly those essential for plant survival. For instance, while we successfully identified *FRI* as a key regulator of flowering time through LoF burden association testing, other well-known regulators such as *FLC* and *FT* were not captured because they did not meet the 5% MAF threshold. That is, the function of genes with few or no natural LoF cannot be studied with this kind of analysis. This limitation is particularly relevant for LoF-intolerant genes, which are often involved in critical biological processes, exhibit low LoF frequencies, and are consequently underrepresented in LoF association testings. These findings align with the recent report by Spence et al. (2024), which highlights that LoF burden tests often fail to prioritize genes based on their importance to a trait. This is because the most trait-critical genes are typically highly constrained and have the lowest LoF variant frequencies. Additionally, Spence et al. noted that gene length significantly influences the power of LoF burden tests, consistent with our observation that coding sequence length is positively correlated with the number of accumulated LoF variants in Arabidopsis. On the other hand, this underscores the importance of accurate LoF variant calling and the aggregation of all LoF variants, as these steps can substantially increase the LoF frequencies of genes of interest, improving their representation in association studies.

## Conclusion and Future Directions

In this study, we explored the characteristics and functional significance of genes tolerant of accumulating putative loss-of-function alleles. Although these genes are not essential for plant survival, they often play specialized roles in processes such as defense and environmental adaptation. Their functions, however, remain underexplored due to functional redundancy and their non-essential nature. To address this, we constructed a comprehensive LoF allele matrix using available whole-genome sequencing data, detailing the functional status of every gene across all accessions represented in the 1001 Genomes Project (**Supplemental Table 1**), in the hope that this LoF matrix will serve as a valuable resource for advancing Arabidopsis genetic studies. Building upon a previous insightful study (Xu et al. 2019) on Arabidopsis LoF variation, our work further incorporates structural variants – an important but sometimes underrepresented source of gene disruption – into LoF calling, and consolidates all LoF variants into gene-level LoF burdens to overcome allelic heterogeneity. We then applied a function-based burden test approach, leveraging gene expression and flowering time as phenotypes to evaluate its effectiveness. Our LoF burden tests successfully identified associations between gene pairs, including those within well-characterized flowering time networks and previously uncharacterized genes, offering insights into both previously undescribed and well-established functional connections.

Future directions for this research include exploring how genes within an organism coordinate to fine-tune complex pathways and traits. For example, can certain pathways or traits be optimized through coordinated changes in gene dispensability? Are there instances where multiple genes co-lose to confer adaptive advantages, and how do these patterns influence fitness across different environmental conditions? Another key question is how dispensable genes contribute to phenotypic plasticity – do they enable rapid adaptation to abiotic stresses such as drought or extreme temperatures? How do they interact with epigenomic marks or regulatory networks to mediate gene expression changes in response to environmental cues? Additionally, what role does genetic redundancy play in buffering mutations and enabling gene loss without detrimental effects on the plant? For applied research, understanding how LoF variants affect agriculturally relevant traits could have significant implications for plant breeding by accelerating the discovery of targets for CRISPR-mediated knockouts. Could we manipulate dispensable genes or create synthetic LoF alleles to improve stress resilience, yield, or nutrient use efficiency? These questions not only provide exciting avenues for future exploration but also underline the importance of studying LoF variants within an integrated framework that connects evolutionary biology, functional genomics, and agricultural applications.

## Supporting information

Summary of LoF burden - flowering time associations

Summary of significant LoF burden - expression associations

Binary collapsed LoF allele matrix for 1,113 accessions and 27,416 genes

## Acknowledgments

We thank members of Monroe Lab for valuable discussions of this work. This work was supported by FFAR grant ICRC20-0000000014 and UC Davis Jastro-Shields Graduate Research Award. Research was conducted at the University of California Davis, which is located on land that was the home of the Patwin people for thousands of years.

## Data and code availability

Figures, supplemental data and code for this research are located at: https://github.com/KehanZhao/ArabidopsisLoF.

## Supplemental Materials

### Supplemental tables

Table S1. Binary collapsed LoF allele matrix for 1,113 accessions and 27,416 genes

Table S2. Summary of significant LoF burden - expression associations

Table S3. Summary of LoF burden - flowering time associations

### Supplemental figures

**Figure S1.**
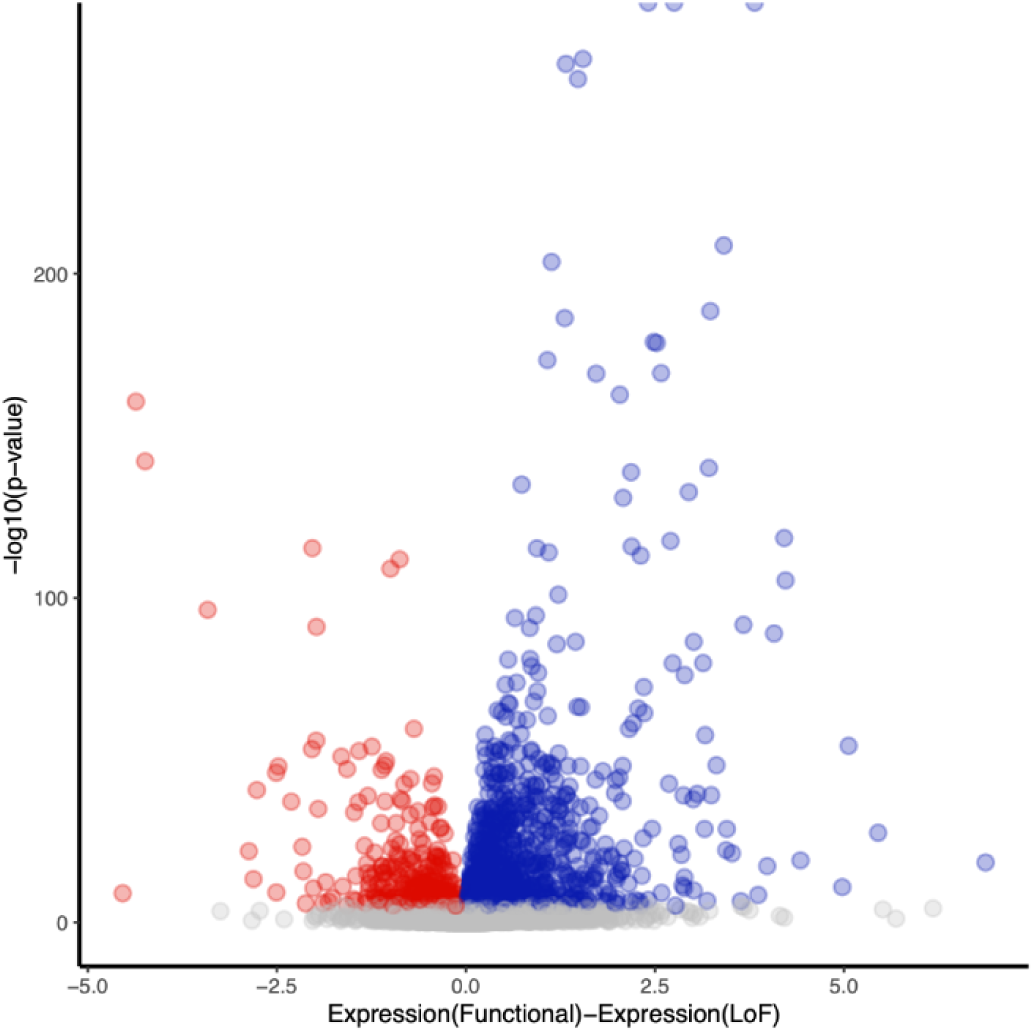
Predicted LoF (pLoF) Alleles Display Generally Lower Expression Level Than Their Functional Counterparts. Each point represents a t-test between the expression level of pLoF alleles and functional alleles of a gene. Blue: functional alleles have higher expression than pLoF alleles; red: pLoF alleles have higher expression than functional alleles; grey: not significant in t-test after multiple testing correction.

**Figure S2.**
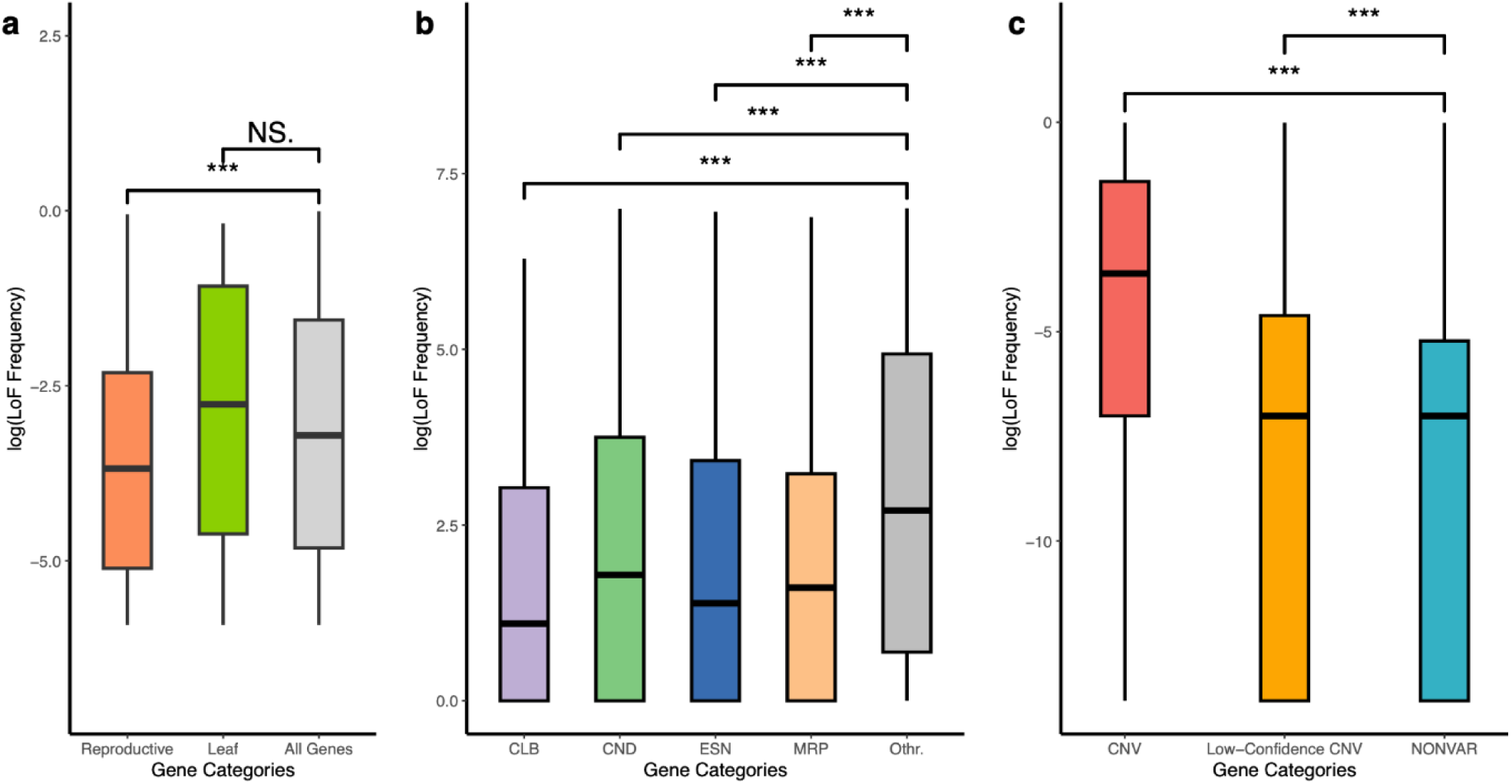
LoF Frequency in Different Gene Categories. a) Box plots of gene LoF frequency of genes highly expressed in leaf tissues, reproductive tissues, and all genes, respectively. Genes highly expressed in one type of tissue is defined as at least twice average transcript level than other tissue types. Leaf tissues include vascular leaf, cauliπe leaf, and rosette leaf. Reproductive tissues include fruit, seed, embryo, flower, silique. Genes highly expressed in reproductive tissues display significantly lower LoF frequency, b) Boxplots of LoF frequency of genes according to their essentiality categories based on Lloyd and Meinke 2012. Genes are categorized based on their mutant phenotypes. CLB: cellular and biochemical; CND: conditional; ESN: essential; MRP: morphological, c) Boxplots of gene LoF frequency of Copy Number Variation (CNV) genes and NONVAR-genes (genes that did not overlap with any CNVs). NONVAR-genes show significantly lower LoF frequency than CNV-genes

**Figure S3.**
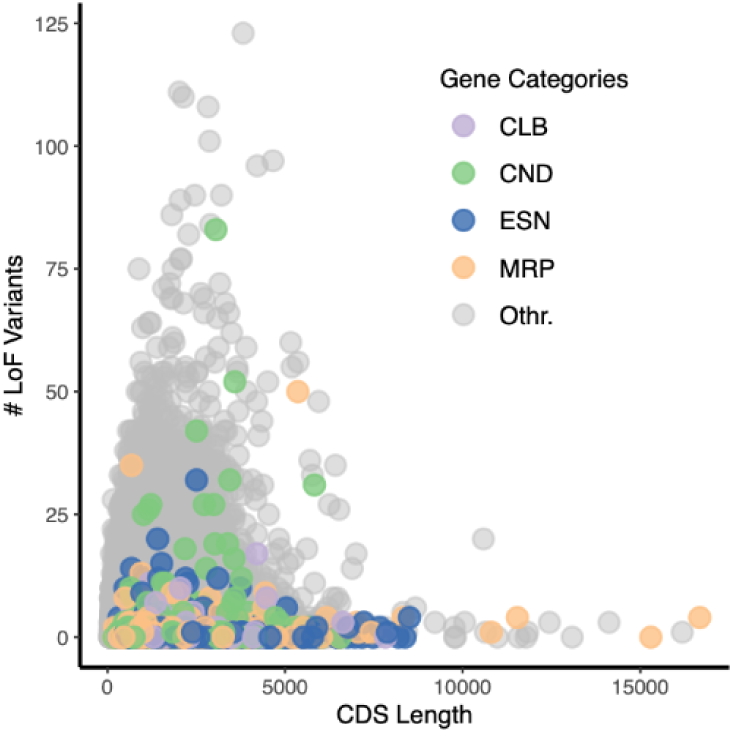
Longer Genes Accumulates More LoF Mutations. A positive correlation was observed between gene CDS length and the number of LoF variants accumulated. Data points were colored by gene categories based on their mutant phenotypes from Lloyd and Meinke 2012. CLB: cellular and biochemical; CND: conditional; ESN; essential; MRP: morphological.

**Figure S4.**
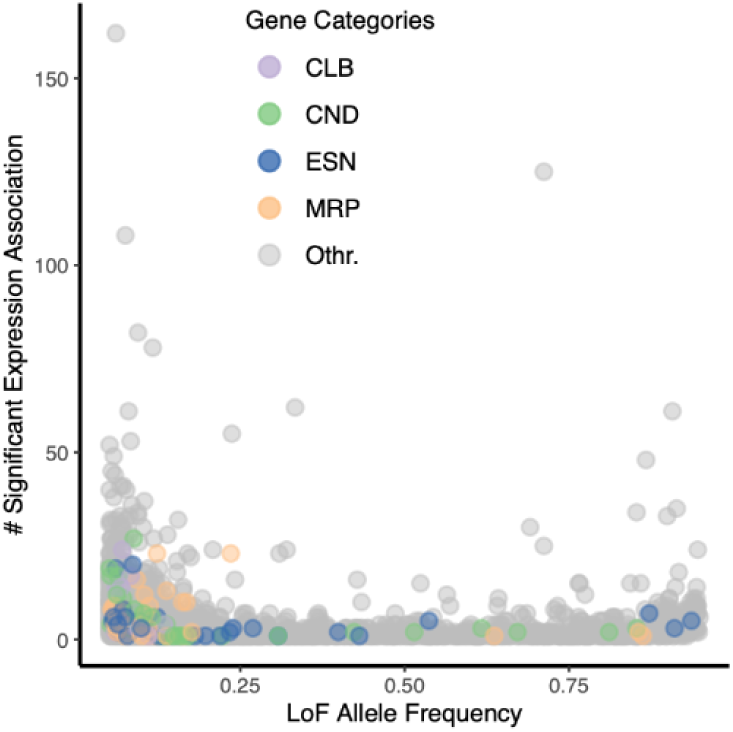
Rare Alleles Show More Expression Associations in LoF Burden Tests. Data points were colored by gene categories based on their mutant phenotypes from Lloyd and Meinke 2012. CLB: cellular and biochemical; CND: conditional; ESN: essential; MRP: morphological.

**Figure S5.**
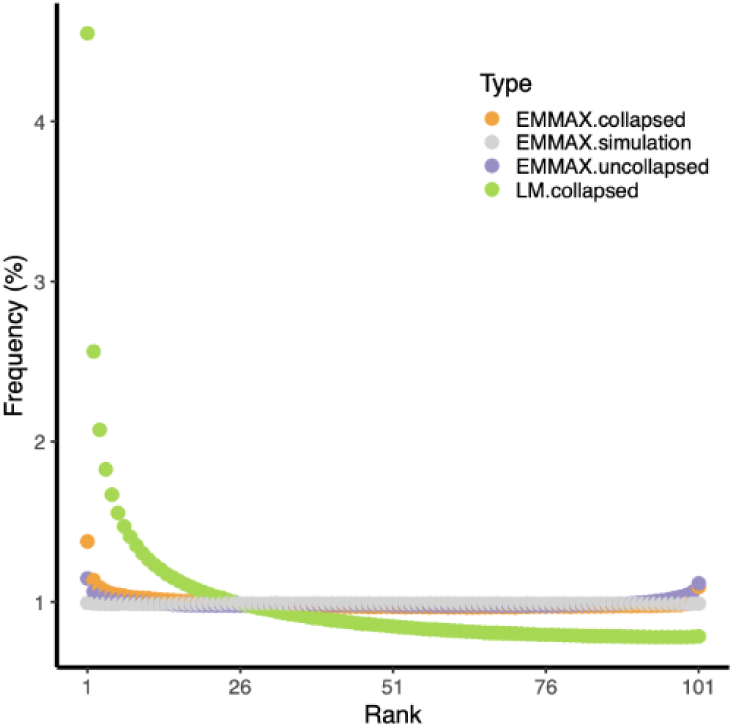
Rank Distribution of LoF-Expression Associations against Simulations Using Different Approaches. Orange: LoF burden tests using EMMAX; Purple: LoF association testing without aggregating LoF burdens; Green: LoF burden tests using a general linear model; Grey: Simulated LoF burden tests using EMMAX by permuting LoF and functional alleles.

**Figure S6.**
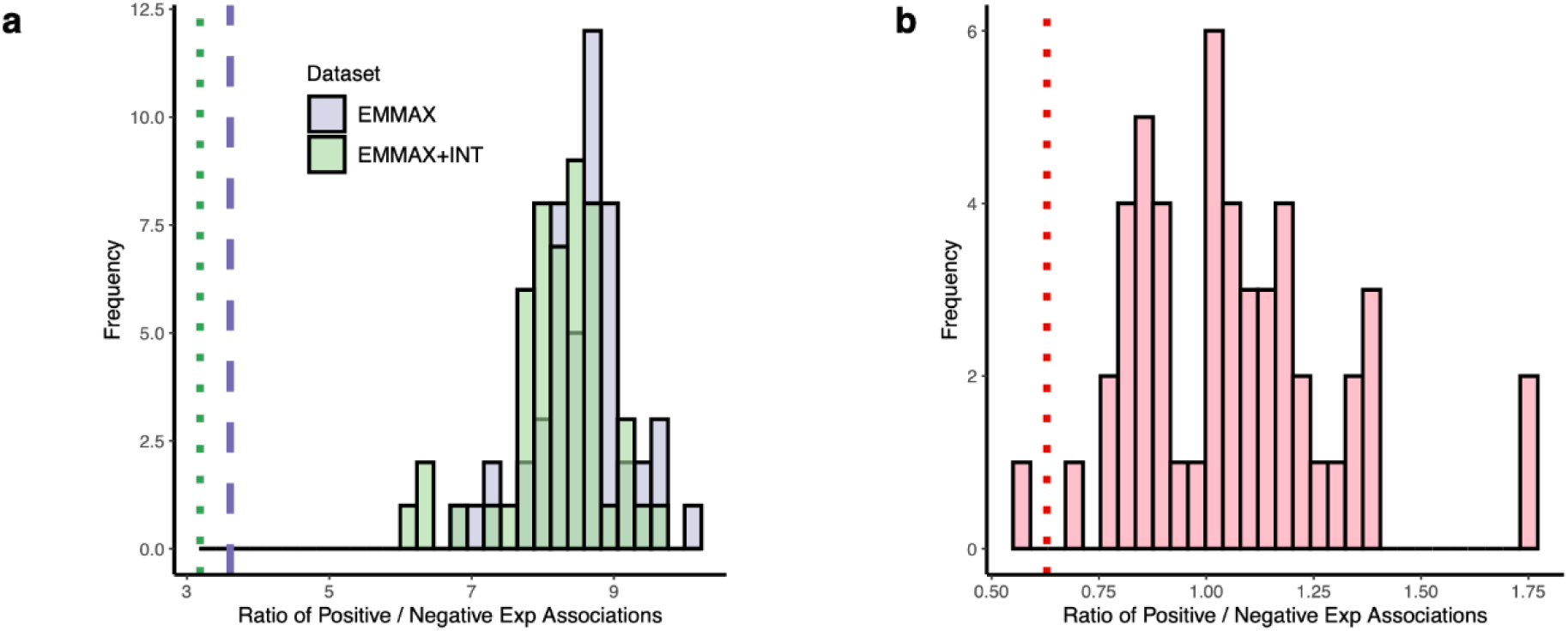
Skewed Expression Data Drives Bias Toward Positive LoF-Expression Associations. a) Histogram of positive-to-negative association ratios from 50 simulated genome-wide LoF burden tests, comparing results before and after inverse normal transformation (INT) of expression data. The purple dashed line indicates the observed positive-to-negative association ratio in the empirical dataset while the green dotted line represents the ratio after INT (∼3.2). b) Histogram of positive-to-negative association ratios from 50 simulated genome-wide LoF burden tests after removing non-normally distributed expression genes. The red dashed line marks the observed positive-to-negative association ratio in the empirical dataset after filtering (∼0.63). Other filtering criteria: 0.05 < LoF allele frequency < 0.95; FDR-corrected p <0.05.

